# Oral administration of GnRH via a cricket vehicle stimulates spermiation in tiger salamanders (*Ambystoma tigrinum*)

**DOI:** 10.1101/2023.08.01.551446

**Authors:** Devin M. Chen, Li-Dunn Chen, Carrie K. Kouba, Nucharin Songsasen, Terri L. Roth, Peter J. Allen, Andrew J. Kouba

**Author notes:** Corresponding author (CKK).

## Abstract

More than 50% of caudates are threatened with extinction and are in need of *ex-situ* breeding programs to support conservation efforts and species recovery. Unfortunately, many salamander populations under human care can experience reproductive failure, primarily due to missing environmental cues necessary for breeding. Assisted reproductive technologies (ARTs) are a useful suite of techniques for overcoming or bypassing these missing environmental cues to promote breeding. Exogenous hormones are used to stimulate natural breeding behaviors or gamete expression for *in-vitro* fertilization or biobanking and are typically administered intramuscularly in caudates. While effective, intramuscular injection is risky to perform in smaller-bodied animals, resulting in health and welfare risks. This research investigated the spermiation response to hormone administration through a non-invasive oral route using the tiger salamander (*Ambystoma tigrinum*) as a model species. Male salamanders were randomly rotated six weeks apart through four treatments (n = 11 males/treatment) in which animals received a resolving dose of gonadotropin-releasing hormone (GnRH) as follows: (1) Prime-Only (0.0 µg/g); (2) Low (0.25 µg/g); (3) Medium (1.0 µg/g); and (4) High (2.0 µg/g). All males were given a GnRH priming dose (0.25 µg/g) 24 hours prior to the resolving dose. Exogenous hormone was delivered inside of a cricket (*Gryllodes sigillatus*) that was presented as a food item by tweezers. Sperm samples were collected at 1, 3, 6, 9, 12, and 24 hours after the resolving dose and analyzed for quantity and quality. For all treatments, sperm concentration was produced in an episodic pattern over time. The Prime-Only treatment had a lower (p < 0.05) percent of sperm exhibiting normal morphology compared to treatments utilizing a resolving dose of GnRH. Overall, oral administration of GnRH is a feasible route of inducing spermiation in salamanders, yielding sperm of sufficient quantity and quality for *in-vitro* fertilization and biobanking efforts.

## Introduction

Amphibians are experiencing some of the highest rates of extinction of all vertebrates, primarily due to habitat destruction, disease, climate change, and pollution [1], resulting in the establishment of conservation breeding programs in zoos as a hedge against extinction [2]. While 41% of all amphibian species are listed as threatened, about 51% of the 805 known caudate species are in decline making them the most at-risk Order within the Class Amphibia [1,3]. For many amphibians, control of reproduction in captivity is essential for sustainability and can be achieved through the manipulation of environmental parameters such as photoperiod, temperature, or humidity [4]. Even under such circumstances, many of the threatened species in human care do not reproduce well and require the use of exogenous hormone administration for stimulating natural reproductive behaviors and gamete production. The use of exogenous hormone therapy for reproductive management in captive breeding programs has been applied to a variety of wildlife taxa including mammals [5-8], birds [9-10], reptiles [11], fishes [12-13], and amphibians [14-15]. One exogenous hormone commonly used in amphibian reproductive therapies is gonadotropin-releasing hormone (GnRH) [16-19]. GnRH is an evolutionarily conserved neurohormone that works via the hypothalamic-pituitary-gonadal (HPG) axis by binding to the pituitary to signal the release of luteinizing hormone and follicle stimulating hormone, which regulate important reproductive functions such as breeding behaviors, steroidogenesis, gametogenesis, spermiation, and ovulation [20].

The use of exogenous hormone therapy for managing caudate reproduction is relatively new compared to several decades of research for anuran species. For reproductive management, anuran species have traditionally been given GnRH through intraperitoneal, intramuscular, dorsal lymph sac, and subcutaneous injections, with the intraperitoneal method being used most frequently [21-24]. In comparison, the attempts to administer GnRH to caudates have generally been via epaxial intramuscular injections [19,25-27]. Many caudate species can be challenging to administer hormones to, not only because of their size (many below 10 grams), but also their quick movements during handling. This makes injections particularly risky (e.g., organ puncturing) for small, slender-bodied, and mucous-producing species, resulting in a need for less invasive methods of hormone delivery.

Thus far, a handful of alternative modes of hormone administration have been explored with some success in anuran species. Frogs and toads have more cutaneous vascularization and granular skin structure in the ventral pelvic region making them more efficient at absorbing aqueous solutions than smooth-skinned salamanders [28]. As such, GnRH mixed with dimethyl sulfoxide (DMSO) applied dermally via the ventral side of American toads (*Anaxyrus americanus*) and Gulf Coast toads (*Incilius valliceps*), produced a spermiation response in more than 70% of individuals [29]. However, the DMSO appeared to cause a red skin irritation in the animals and was not subsequently adopted as an alternative method of hormone delivery. More simply, ovulation in African clawed frogs (*Xenopus laevis*) resulted from the passive absorption of human chorionic gonadotropin (hCG) via a water bath [30]. Similarly, dermal application of GnRH to Texas blind salamanders (*Eurycea rathbuni*) stimulated reproduction [31], which is likely due to the porous nature of their skin as an obligate aquatic species. However, dermal administration of hormones in terrestrial caudates has not been examined.

Nasal and oral delivery of exogenous hormones are alternative routes that have been successful in inducing gamete expression in other taxa and in anuran amphibians. Intranasal administration of GnRH was tested in Fowler’s toads (*Anaxyrus fowleri*) and induced spermiation in 93% of the males treated, with high motility and sufficient concentrations of sperm for cryopreservation and *in-vitro* fertilization [17]. Overflow of hormone was swallowed in the nasal treatment suggesting that oral hormone administration could be effective [17]. Subsequently, GnRH administered directly into the mouths of Fowler’s toads, without a vehicle, was observed to induce spermiation [32], though not as effectively as nasal administration possibly due to stomach proteases breaking down the hormone. However, GnRH given via a mealworm vehicle in American toads caused only one-third of males to spermiate [29]. Effective delivery of exogenous hormones through the gastrointestinal tract may depend on the type of vehicle and to what extent the vehicle could be digested. Endogenous GnRH has a short half-life of 2-4 min and is broken down by peptidases, though the analogues used in most research have a half-life of a few hours [33]. Although both oral and nasal administration of exogenous hormones for stimulating spermiation have been tested in anurans, the use of oral routes of hormone delivery in salamander species is lacking.

The purpose of this study was to examine whether oral administration of GnRH, via a cricket (*Gryllodes sigillatus*) vehicle, elicits spermiation in eastern tiger salamanders (*Ambystoma tigrinum*), a model species for the development of assisted reproductive technologies (ARTs) in caudates. Crickets were chosen as a vehicle because they digest more readily compared to mealworms and are broadly available. The specific aims of the study were to test the impact of using an insect vehicle for GnRH on: (1) the percent of males producing sperm; (2) sperm quantity [concentration, total sperm cells]; and (3) sperm quality [motility parameters, viability, morphology] over time in a dose-dependent manner. We hypothesized that oral administration of GnRH via a cricket would cause reliable spermiation due to the ability of the hormone to easily exit the cricket vehicle into the salamander’s system. We further predicted that the largest dose treatment of GnRH would result in the highest number of sperm producers, with the greatest sperm quantity, and significantly better sperm quality indices. Oral hormone administration could advance breeding efforts and increase genetic diversity by mitigating health risks and improving the welfare of caudates in *ex-situ* conservation breeding programs. Additionally, the ease of hormone delivery and low-risk approach may expand accessibility to ART procedures for breeding practitioners.

## Materials and methods

### Experimental animals

Captive-bred, sexually mature male eastern tiger salamanders (mean mass = 60.5 ± 11.1 g) were housed in 30 x 46 x 66 cm enclosures, with 5 cm of coconut fiber as bedding, in groups of 2–6 animals [27]. Salamanders were kept on a 12-hour light cycle and provided with water baths as well as clay and polyvinyl chloride hides and were fed a diet of crickets (*G. sigillatu**s***), mealworms (*Tenebrio molitor*), and Dubia roaches (*Blaptica dubia*) dusted with calcium supplement (Zoo Med Laboratories, Inc., CA, USA) and vitamin mix (Supervite, Repashy Ventures Inc., CA, USA). Previous research with these study animals found that seasonality did not have an effect on spermic response [26], but hormone was not administered between December and February (natural breeding season) for this study as a precaution. During experimentation, males were kept individually in 28 x 15 x 13 cm plastic tubs with 200 mL of water for 48 hours. Approximately 30 minutes before receiving a priming dose of GnRH, salamanders were placed into the holding container for acclimation and were returned to their standard enclosures one hour after the final sperm collection. Salamanders were individually identifiable based on unique skin patterns and chin spots. All research was approved under IACUC #20-160 at Mississippi State University.

### Exogenous hormone administration

Male tiger salamanders (n = 11 males/treatment) were administered one of four hormone treatments that consisted of a priming dose and resolving dose of GnRH (Prime-Only, Low, Medium, or High), as shown in Fig 1. Salamanders were rotated through the hormone treatments at least six weeks apart to ensure hormone clearance from any prior treatment, as previously recommended [26,34] The GnRH analogue (Ala6, des-Gly10 ethylamide LHRH; Sigma-Aldrich® catalog #L4513) was reconstituted to a concentration of 1 µg/µL or 5 µg/µL (depending on volume of final dose to be given that could fit within a cricket) in phosphate buffered saline (PBS) and administered via an insect vehicle in which the body of a G. *sigillatus* was filled with hormone, via pipette, following removal of the cricket’s head by hand and clearing the insides by pipette aspiration (Fig 2). All males were given a priming dose of 0.25 µg/g body weight (BW) GnRH in the cricket vehicle. The resolving dose of GnRH for each treatment was similarly delivered 24 hours after the priming dose as follows: Prime-Only (PBS given as resolving dose; 0.0 µg/g BW), Low (0.25 µg/g BW), Medium (1.0 µg/g BW), or High (2.0 µg/g BW). Crickets with specific hormone treatments were presented via tweezers to the salamanders for feeding (Fig 2). Typically, salamanders ate their crickets in one swift motion and swallowed the crickets whole within two minutes, minimizing any hormone loss. The salamanders in the study are regularly fed crickets in this manner as part of their usual diet.

**Fig 1.**
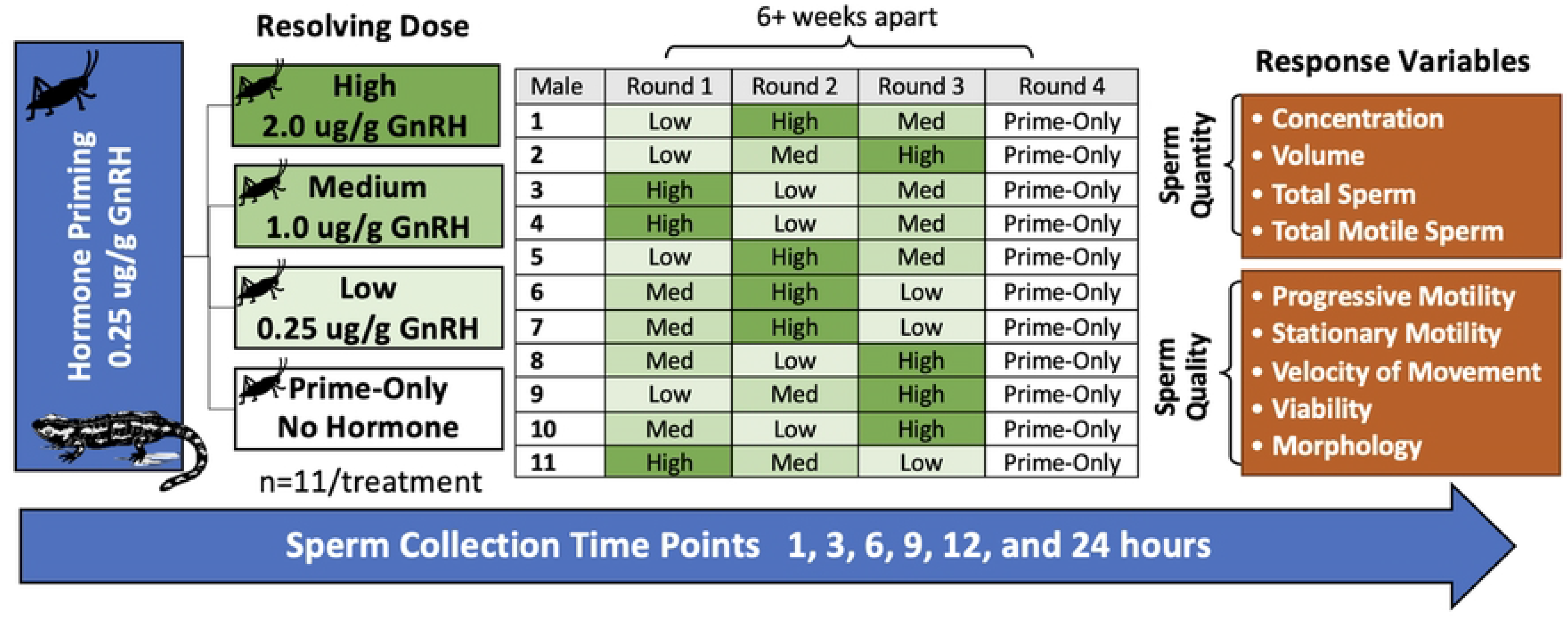
Experimental design for studying the effect of orally-delivered GnRH on tiger salamander sperm production. Male tiger salamanders (*Ambystoma tigrinum*) (n = 11) were cycled through Prime-Only, Low, Medium, and High GnRH oral administration hormone treatments. Response variables included both sperm quantity and quality measures and parameters were measured at 1, 3, 6, 9, 12, and 24 hours after the resolving dose.

**Fig 2.**
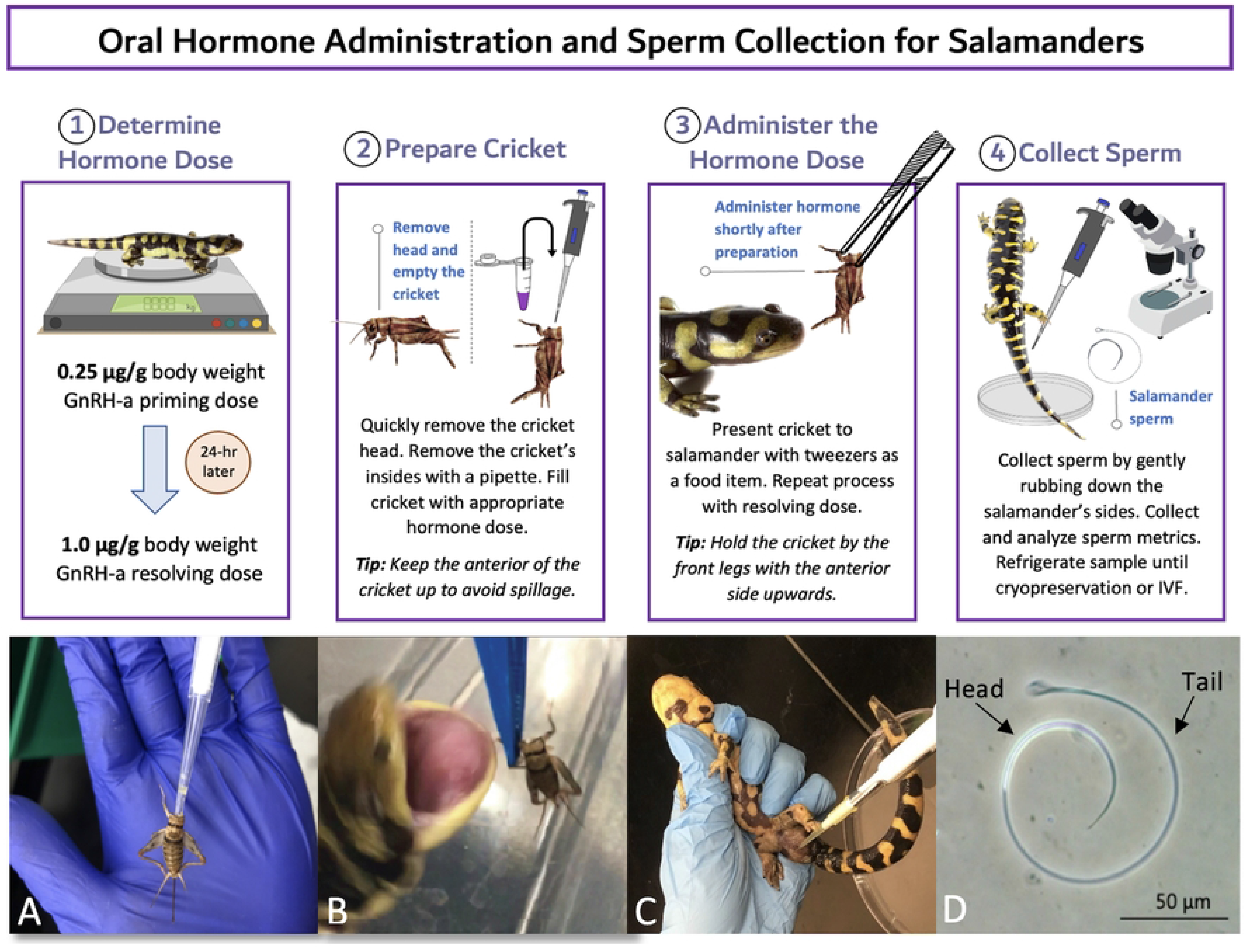
Diagram outlining the protocol for delivering hormones orally to caudates. Below: A) Cricket being emptied using a pipette; B) Tiger salamander (*Ambystoma tigrinum*) being fed a treatment cricket; C) Collection of spermic material from the cloaca of a male *A. tigrinum*; and D) *A. tigrinum* sperm cell under 200x magnification.

### Sperm collection and analysis

All males were checked for sperm production prior to administration of the priming dose, and then at 1, 3, 6, 9, 12, and 24 hours after the resolving dose was given. Sperm was expressed by lightly tapping the cloaca of the salamander for 10 seconds then massaging downwards on the lateral sides of the animal for another 10 seconds, and finally applying light pressure above the pelvis on each side of the animal. Sperm was excreted as either milt (viscous, white material with a high concentration of sperm) or spermic urine (non-viscous, clear material with a lower concentration of sperm), as previously described [26-27]. Sperm was collected via a pipette from the cloaca and immediately analyzed for several sperm quality parameters including motility, morphology, and viability. Motility parameters included progressive motility (PM; sperm actively rotating in a circular motion), stationary motility (SM; presence of undulating tail membrane), and total motility was calculated (TM = PM + SM). The sperm ‘velocity of movement’ (observer scale from 1-5, where 5 indicates maximal circular rotation and 1 indicates minimal) was also recorded. Sperm morphology was categorized as either normal, broken head, broken tail, or broken head plus tail [35]. The proportion of motile cells and morphology parameters were counted out of 100 random sperm cells under a 20x objective lens using a phase contrast microscope (Olympus CX41). Viability was measured using an Invitrogen™ Live/Dead stain containing SYBR14/propidium iodide where live (green) and dead (red) sperm cells were counted out of 100 random cells on the slide using an Olympus CX41 with fluorescent attachment (Fig 3). Following sperm quality evaluation, sperm quantity parameters were measured using a Neubauer haemocytometer including concentration (sperm/mL), volume, total number of sperm cells (volume x concentration), and total number of motile sperm cells (total number of sperm cells x TM). The number of overall spermic responders was additionally compared between treatments. A spermic response occurred if males were not releasing countable sperm before a hormone treatment but subsequently released sperm in the form of spermic urine or milt.

**Fig 3.**
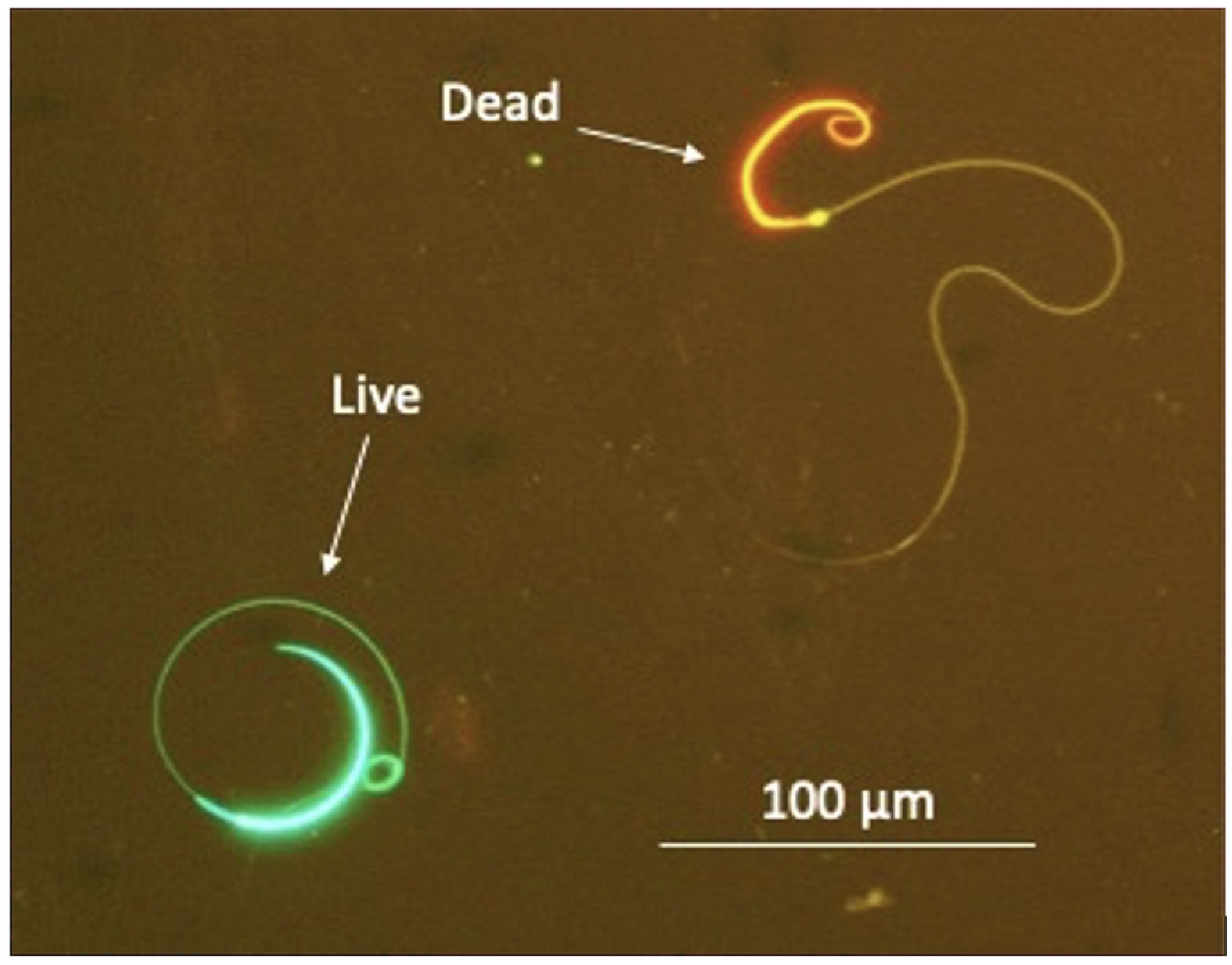
Live/Dead stain of salamander sperm using SYBR14/propidium iodide to indicate sperm sample viability. Damaged cells fluoresce red whereas intact cells fluoresce green.

### Case study

We further tested the Medium oral dose of GnRH (0.25 µg/g prime + 1.0 µg/g resolving dose) in a smaller (mean mass = 13.0 ± 1.2 g), slender-bodied plethodontid species, the arboreal salamander (*Aneides lugubris*). This species can be difficult to deliver hormones to through an intramuscular injection and serves as a good case study for oral hormone administration in caudates. Four captive male arboreal salamanders were administered GnRH using small, 1/8” *G. sigillatus* crickets as described in Fig 2. Males were checked for sperm at 1, 3, 5, 7, and 24 hours post-resolving dose.

### Statistical analysis

Data were checked for normality using a Shapiro-Wilk test as well as histogram visualization, whereas homogeneity of variance was checked with a Levene’s test. The package ‘fitdistrplus’ was employed to find data families. A Generalized Additive Model for Location, Scale and Shape (GAMLSS) was used for regression modeling, where hormone treatment was the fixed effect and individual was the random effect. Time and a treatment by time interaction were also tested. Either a beta, beta zero-inflated, generalized gamma, generalized inverse gamma, Gumbel, normal, double Poisson, or Poisson family was used depending on the distribution of the response variable. Model fit was checked using QQ plots as well as testing the normality of the residuals. The number of responders to each treatment was compared using two-tailed Fisher’s exact tests. Data are shown as mean ± SEM, with the alpha value set to 0.05 for significance testing. Data were analyzed in the statistical software, R [36]. Due to the small sample size, the case study results were evaluated using descriptive statistics only.

## Results

### Number of spermic responders

The hormone treatments induced spermiation in 81% (9/11), 64% (7/11), 73% (8/11), and 81% (9/11) of males in response to the Prime-Only, Low, Medium, and High treatments of GnRH, respectively, at a minimum of one sampling point. The number of responders between each treatment was similar (p > 0.05), thus the Prime-Only treatment stimulated a majority of animals to produce sperm and the addition of a second GnRH treatment did not induce spermiation in additional animals. This indicates a single low dose of hormone can be given orally to collect sperm from a similar number of animals as seen utilizing an additional resolving dose of GnRH.

### Sperm parameters within a treatment averaged across time

The means and ranges of the spermic response parameters are shown in Table 1. Caudate sperm is expressed in three sample types as follows: spermic urine (below 1 x 10^6^ sperm/mL), milt (above 1 x 10^6^ sperm/mL), and spermatophores (not observed in this experiment). There was a total of 128 spermic samples collected from the eleven males across the six sampling points for all four treatments; there was an average of 31 spermic samples per treatment. Overall, the proportion of samples collected was heavily weighted (p < 0.05) toward spermic urine (samples below 1 x 10^6^ sperm/mL) compared to milt (samples above 1 x 10^6^ sperm/mL) collections (Fig 4A). There was no difference (p > 0.05) in sperm concentration within sample type between each treatment. Sperm concentrations ranged from x 10^3^ to 10^7^ (Fig 4B), with individuals typically exhibiting an oscillatory sperm production pattern switching between spermic urine and milt production. Overall, sperm concentration was similar (p > 0.05) for the Prime-Only (1.5 ± 0.5 x 10^6^ sperm/mL), Low (0.9 ± 0.3 x 10^6^ sperm/mL), Medium (2.0 ± 0.7 x 10^6^ sperm/mL), and High (0.7 ± 0.3 x 10^6^ sperm/mL) treatments (Table 1). When adjusting for volume of spermic urine (98 ± 17 µL) and milt (37 ± 13 µL) to calculate average total sperm cells produced within each treatment, we found no difference (p > 0.05) between the Prime-Only (0.45 ± 0.14 x 10^5^), Low (0.34 ± 0.11 x 10^5^), Medium (0.37 ± 0.14 x 10^5^), or High (0.32 ± 0.11 x 10^5^) GnRH treatments (Table 1). Similarly, the total number of motile sperm cells between the Prime-Only (0.30 ± 0.88 x 10^5^), Low (0.26 ± 0.85 x 10^5^), Medium (0.30 ± 0.13 x 10^5^), and High (0.21 ± 0.68 x 10^5^) treatments were similar (p > 0.05).

**Fig 4.**
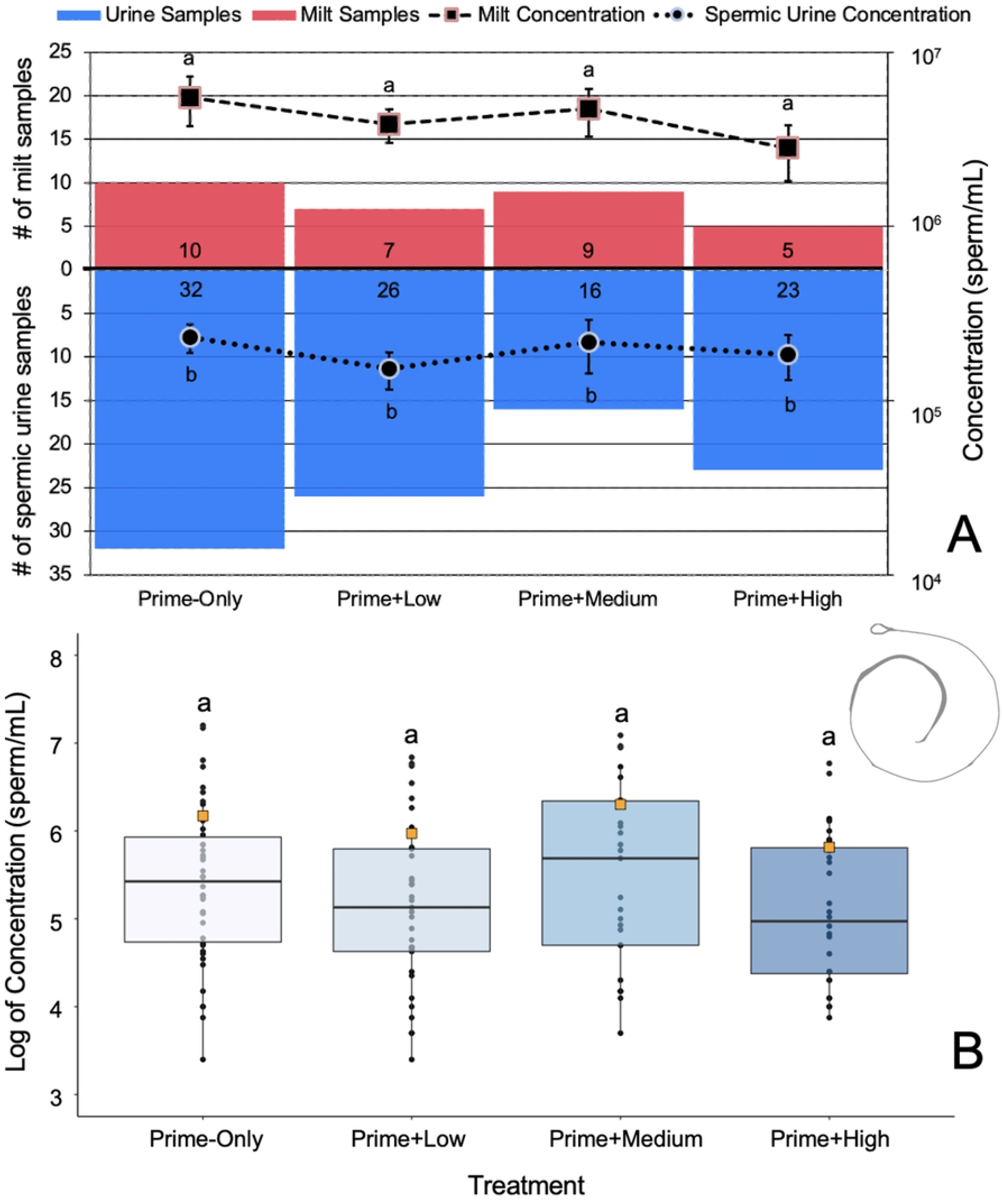
Comparison of sperm sample concentrations between treatments. (A) Number of spermic urine (blue) versus milt (red) samples for each hormone treatment. The dashed line highlights the average milt concentration between each treatment, whereas the dotted line highlights the average spermic urine concentration. Letters indicate significant differences within and between sample types. (B) Concentration of sperm samples averaged across sample type and time points for each hormone treatment. Boxes represent the interquartile range, whiskers show the range of the data, the horizontal bars indicate the median, the orange squares represent the mean, and different letters indicate significant differences between treatments (p < 0.05).

**Table 1.**
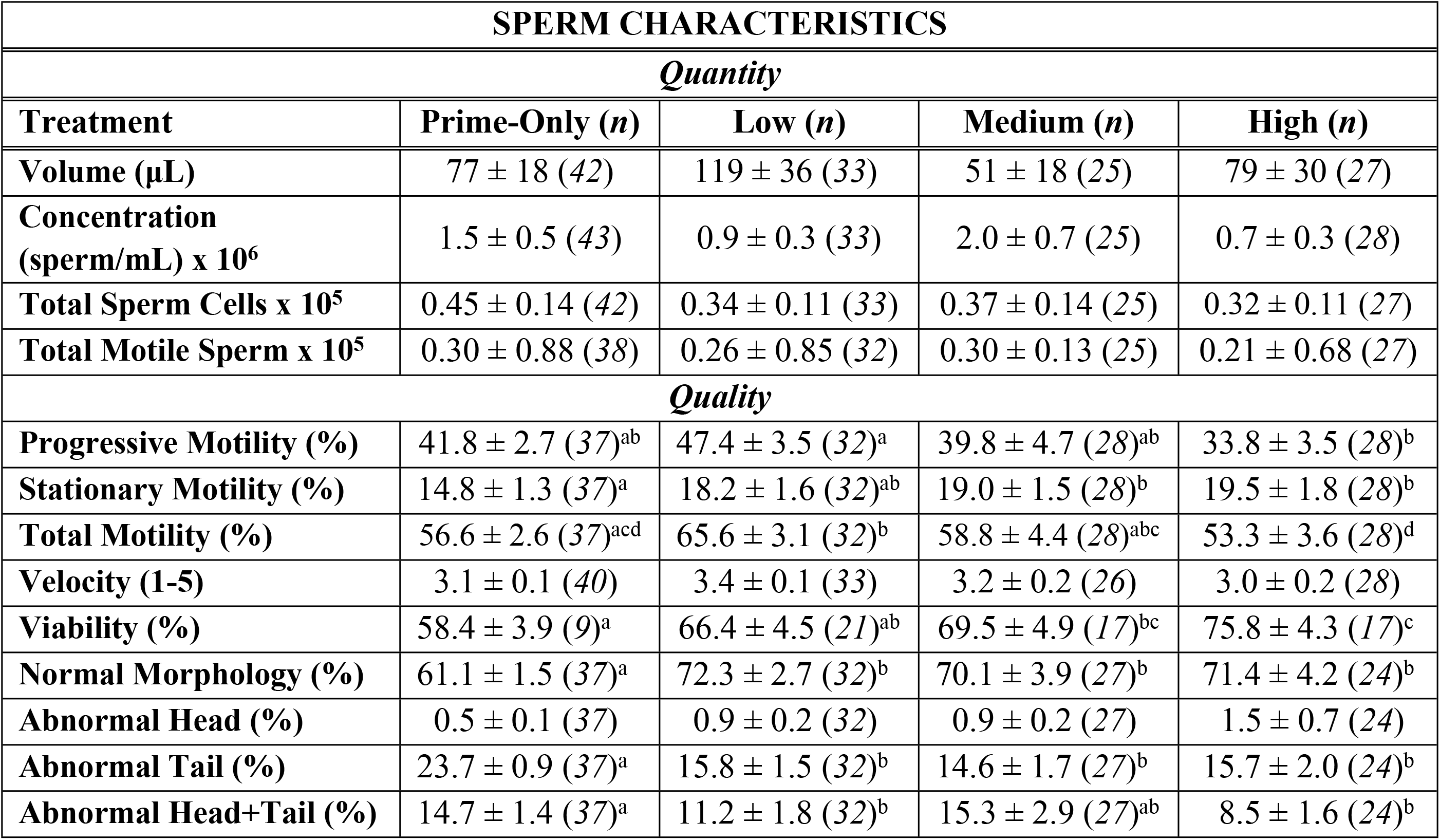
Sperm characteristics compared between oral hormone treatments. A comparison of mean ± SEM sperm metrics among Prime-Only, Low, Medium, and High oral GnRH treatments in male tiger salamanders (*Ambystoma tigrinum*) (n = 11). Averages of spermic samples are combined across all individual time points for all males within a treatment group. The numbers in parentheses indicate the number of sperm samples used to calculate the metric. Different letters indicate significant differences (p < 0.05).

| SPERM CHARACTERISTICS |  |  |  |  |
| --- | --- | --- | --- | --- |
| *Quantity* |  |  |  |  |
| Treatment | Prime-Only (n) | Low (n) | Medium (n) | High (n) |
| Volume (μL) | $77 \pm 18$ (42) | $119 \pm 36$ (33) | $51 \pm 18$ (25) | $79 \pm 30$ (27) |
| Concentration (sperm/mL) $\times 10^6$ | $1.5 \pm 0.5$ (43) | $0.9 \pm 0.3$ (33) | $2.0 \pm 0.7$ (25) | $0.7 \pm 0.3$ (28) |
| Total Sperm Cells $\times 10^5$ | $0.45 \pm 0.14$ (42) | $0.34 \pm 0.11$ (33) | $0.37 \pm 0.14$ (25) | $0.32 \pm 0.11$ (27) |
| Total Motile Sperm $\times 10^5$ | $0.30 \pm 0.88$ (38) | $0.26 \pm 0.85$ (32) | $0.30 \pm 0.13$ (25) | $0.21 \pm 0.68$ (27) |
| *Quality* |  |  |  |  |
| Progressive Motility (%) | $41.8 \pm 2.7$ (37)<sup>ab</sup> | $47.4 \pm 3.5$ (32)<sup>a</sup> | $39.8 \pm 4.7$ (28)<sup>ab</sup> | $33.8 \pm 3.5$ (28)<sup>b</sup> |
| Stationary Motility (%) | $14.8 \pm 1.3$ (37)<sup>a</sup> | $18.2 \pm 1.6$ (32)<sup>ab</sup> | $19.0 \pm 1.5$ (28)<sup>b</sup> | $19.5 \pm 1.8$ (28)<sup>b</sup> |
| Total Motility (%) | $56.6 \pm 2.6$ (37)<sup>acd</sup> | $65.6 \pm 3.1$ (32)<sup>b</sup> | $58.8 \pm 4.4$ (28)<sup>abc</sup> | $53.3 \pm 3.6$ (28)<sup>d</sup> |
| Velocity (1-5) | $3.1 \pm 0.1$ (40) | $3.4 \pm 0.1$ (33) | $3.2 \pm 0.2$ (26) | $3.0 \pm 0.2$ (28) |
| Viability (%) | $58.4 \pm 3.9$ (9)<sup>a</sup> | $66.4 \pm 4.5$ (21)<sup>ab</sup> | $69.5 \pm 4.9$ (17)<sup>bc</sup> | $75.8 \pm 4.3$ (17)<sup>c</sup> |
| Normal Morphology (%) | $61.1 \pm 1.5$ (37)<sup>a</sup> | $72.3 \pm 2.7$ (32)<sup>b</sup> | $70.1 \pm 3.9$ (27)<sup>b</sup> | $71.4 \pm 4.2$ (24)<sup>b</sup> |
| Abnormal Head (%) | $0.5 \pm 0.1$ (37) | $0.9 \pm 0.2$ (32) | $0.9 \pm 0.2$ (27) | $1.5 \pm 0.7$ (24) |
| Abnormal Tail (%) | $23.7 \pm 0.9$ (37)<sup>a</sup> | $15.8 \pm 1.5$ (32)<sup>b</sup> | $14.6 \pm 1.7$ (27)<sup>b</sup> | $15.7 \pm 2.0$ (24)<sup>b</sup> |
| Abnormal Head+Tail (%) | $14.7 \pm 1.4$ (37)<sup>a</sup> | $11.2 \pm 1.8$ (32)<sup>b</sup> | $15.3 \pm 2.9$ (27)<sup>ab</sup> | $8.5 \pm 1.6$ (24)<sup>b</sup> |

Whereas the Prime-Only treatment produced samples of similar sperm quantity, the additional resolving doses did increase some sperm quality parameters. The Prime-Only treatment (56.6 ± 2.6%) had lower (p < 0.05) total motility compared to the Low GnRH treatment (Fig 5A). However, the High treatment (53.3 ± 3.6%) resulted in lower (p < 0.05) total motility than the Low (65.6 ± 3.1%) and Medium treatments (58.5 ± 4.4%) (Fig 5A), suggesting hormone dosage could be influencing sperm release. The velocity of movement was equivalent (p > 0.05) between the four treatment groups (Fig 5B). However, more viable sperm (p < 0.05) were released when given the High (75.8 ± 4.3%) or Medium (69.5 ± 4.9%) GnRH doses than the Low (66.4 ± 4.5%) dose or Prime-Only (58.4 ± 3.9%), suggesting a dose dependency (Fig 5C). Importantly, a higher proportion (p < 0.05) of morphologically normal sperm were released when animals were given a resolving dose of GnRH (High, Medium, or Low) than if given only a prime dose of GnRH (Fig 5D). More specifically, abnormal sperm tails were more likely (p < 0.05) to occur in Prime-Only samples compared to the other three treatments.

**Fig 5.**
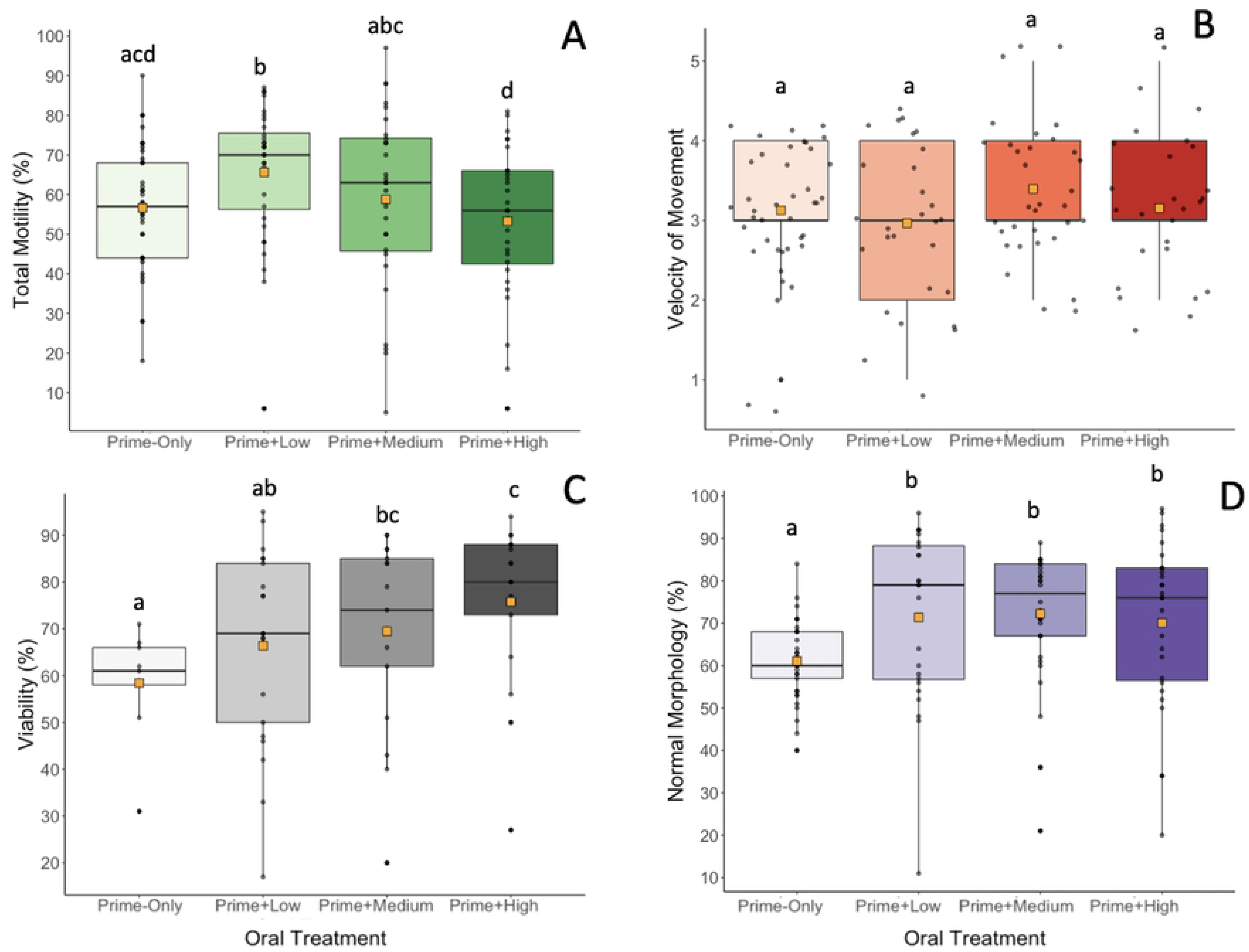
(A) Total motility; (B) velocity of movement; (C) viability; and (D) normal morphology of samples collected from male tiger salamanders (*Ambystoma tigrinum*) comparing Prime-Only, Low, Medium, and High GnRH oral hormone treatments. Boxes represent the interquartile range, whiskers show the range of the data, the horizontal bars indicate the median, the orange squares represent the mean, and different letters indicate significant differences (p < 0.05).

### Individual sperm parameters over time within a treatment

In terms of the sperm quantity and quality over time, both milt and spermic urine samples were produced episodically for all treatments, with tiger salamanders exhibiting an episodic, intervallic sperm expression pattern where a surge of more highly concentrated milt is typically surrounded by periods of lower concentrated spermic urine production (Fig 6A) [26]. We found some treatment by time interactions indicating that the magnitude of sperm production depends on the hormone dose level and collection period (Fig 6A). For example, the highest sperm concentration occurred at 24 hours for the Prime-Only treatment (2.8 ± 1.9 x 10^6^ sperm/mL), 12 hours for the Low treatment (1.4 ± 2.2 x 10^6^ sperm/mL), one hour for the Medium treatment (4.2 ± 6.9 x 10^6^ sperm/mL) and three hours for the High treatment (2.9 ± 2.6 x 10^6^ sperm/mL) (Table 1; Fig 6A). There was no significant difference (p > 0.05) in any of the other measured sperm metrics between time points when averaged and compared across the four treatments (Fig 6B-D).

**Fig 6.**
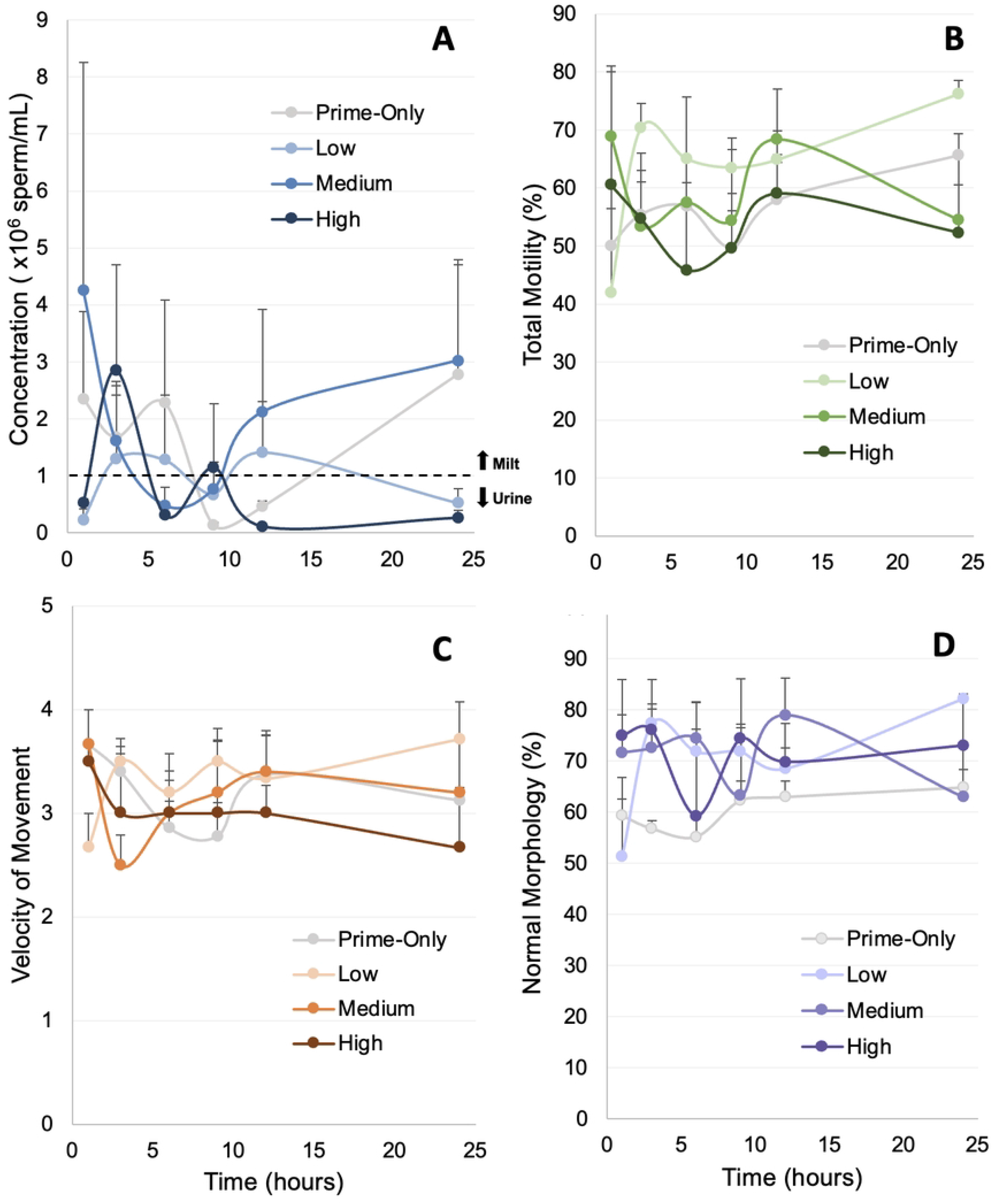
Time course showing the change in mean ± SEM in (A) sperm concentration, (B) total motility, (C) velocity of movement, and (D) normal morphology over time across treatments. Samples were collected at 1, 3, 6, 9, 12, and 24 hours post-resolving dose. Note the episodic surges in sperm concentration.

### Case study

Oral administration of GnRH via a cricket vehicle was transferable to a smaller salamander species, as three out of four male arboreal salamanders produced a spermic response. Sperm concentration was highest at 3- and 5-hours post-resolving dose (Fig 7) compared to the earlier and later sperm collections, while total sperm motility stayed relatively constant over time. The overall average sperm concentration was 1.9 ± 1.2 x 10^6^ sperm/mL, whereas the overall average sperm total motility was 46.3 ± 6.7%. Sperm was of high enough concentration to use for *in-vitro* fertilization, and we were able to cryopreserve several samples following the methods used in tiger salamanders [38].

**Fig 7.**
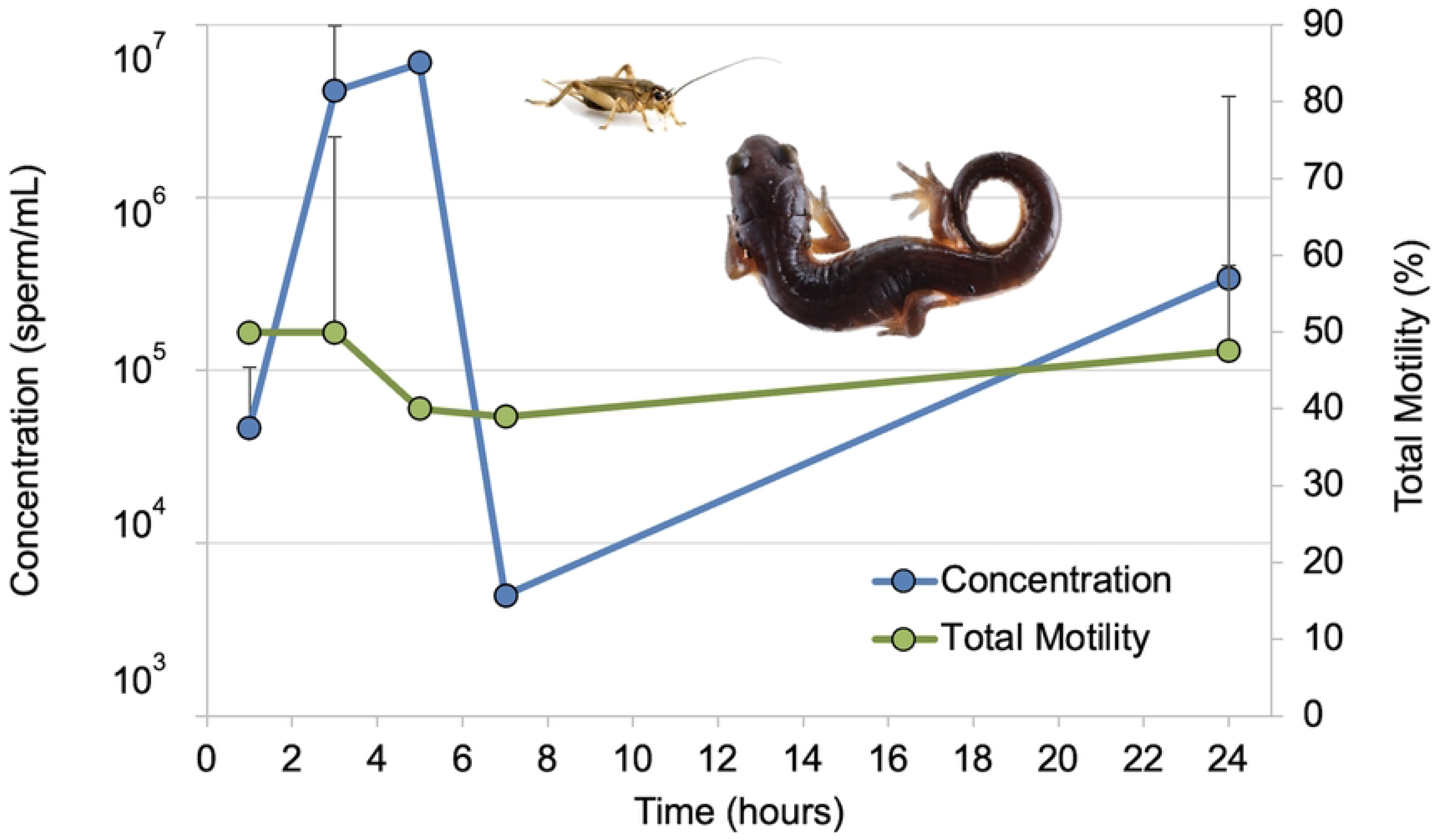
Application of the Medium (0.25 µg/g prime + 1.0 µg/g resolving dose) oral hormone dose in a smaller plethodontid species, *Aneides lugubris*. Time course showing the change in mean sperm concentration and total motility over time. Samples were collected 1, 3, 5, 7, and 24 hours post-resolving dose. Data points with standard deviation bars indicate time points where samples from two individuals were collected. Data points without error bars indicate data were recorded for only one male.

## Discussion

Currently, the most common form of hormone administration for caudates is intramuscular injection, which is risky to perform in small or slender-bodied caudates that have shallow body depths relative to syringe needles (typically 27-31 gauge, 8 mm depth). Hence, reliable and less invasive methods of hormone administration are needed as a growing number of *ex-situ* breeding programs are established for these smaller, threatened caudate species. We demonstrate that oral administration of GnRH through a cricket vehicle is a dependable method to induce spermiation in male tiger salamanders and arboreal salamanders. Not only did a majority of males produce sperm, but the sperm collected over time was of sufficient quantity and quality for use in *in-vitro* fertilization and gamete cryopreservation. We found that although certain sperm quality characteristics were dose dependent, the number of spermic responders and the resulting sperm concentrations were similar for all hormone doses tested.

Previous research [26] found that 73% of male tiger salamanders spermiated in response to injections of GnRH into the epaxial muscle, which is similar to what we observed using oral delivery with a cricket vehicle. However, hormone sequestered into a cricket vehicle produced more than twice the number of spermic responders compared to mealworms injected with 100 µg of GnRH for American toads [29]. The thick, chitinous exoskeleton of the mealworm could be preventing the diffusion and absorption of GnRH into the amphibian circulatory or lymphatic system. In our study, the head of the cricket was removed to insert the hormone (as opposed to a syringe injection into the cricket), which allowed for GnRH to quickly exit into the salamander’s digestive system and be absorbed systemically. Furthermore, the digestive tract of the cricket was removed prior to filling in hormone, removing some digestive enzymes [39] that could break down the hydrolysable peptide bonds of GnRH. While proteolytic enzymes in the salamander digestive tract are also potential agents of hormone breakdown, the cricket exoskeleton and fatty tissue likely helped protect the GnRH. It is also possible that the hormone exited the cricket and reached the GnRH receptors that exist in the lamina propia of the nasal cavity and mucosa that connects to the mouth of the salamander before reaching the stomach [37]. To note, the GnRH analogue implemented in this research lasts on the time scale of hours rather than minutes. An alternative oral approach of pipetting GnRH directly into the mouths of Fowler’s toads (*Anaxyrus fowleri*) has been studied and resulted in an average of 1.8 ± 0.4 x 10^5^ sperm/mL [32], which is lower than the average of 1.9 ± 1.2 x 10^6^ sperm/mL we observed using a cricket vehicle; however, this is a plausible form of oral hormone administration in cases where animals do not willingly eat prey items presented via tweezers.

Oral administration of GnRH resulted in surges of sperm maturation and release similar to what has been seen using intramuscular injections [26]. However, differences in the timing of peak sperm concentrations were observed, which could be due to differences in hormone absorption resulting in the timing of GnRH release, thereby impacting spermiation. In tiger salamanders, intramuscular administration of GnRH resulted in 6.0 ± 1.5 x 10^7^ sperm/mL [26], whereas oral administration resulted in an average of 2.0 ± 3.4 x 10^6^ sperm/mL. Although this is an order of magnitude less, a concentration above 0.5 x 10^6^ sperm/mL is suitable for the fertilization of eggs in this species. For example, 88 viable tiger salamander larvae were recently produced using cryopreserved sperm at a final concentration of 0.5 x 10^6^ sperm/mL [38]. When GnRH was intramuscularly administered to other caudate species, sperm concentrations ranged from 0.3 to 58.0 x 10^6^ sperm/mL (sharp-ribbed newt [39]; eastern hellbender [40]; black-spotted newt [19]; Kweichow newt [19]). While sperm concentration is influenced by species, hormone dose and type, time of collection, and route of administration, the average sperm concentrations produced by both tiger and arboreal salamanders from oral GnRH administration falls within a similar range compared to other caudate species given intramuscular injections.

We found that tiger salamanders treated with GnRH exhibit an episodic pattern of sperm production where periods of highly concentrated milt were released followed by periods of spermic urine, with milt typically resurging several hours later. This bi- or tri-modal episodic pattern in milt production differs from anuran species that have a unimodal distribution in sperm concentration [23,41]. The episodic pattern in salamander sperm production is a function of when milt versus spermic urine is released and is likely affected by their natural reproductive strategy of producing highly concentrated sperm masses packaged within an ampulla or spermatophore and depositing these capsules over time in an aquatic or moist environment [42]. Whereas sperm concentration of milt was an order of magnitude higher than that of spermic urine across all treatments, we chose to combine the values across all time points to acquire a mean sperm concentration for the entire collection period, which is typically the procedure when collecting sperm for storage or transport. When milt and spermic urine were combined, there was no difference in total sperm output between the treatments over time due to the high variability in concentration between sperm types. The mixing of milt with spermic urine can effectively serve as a method of naturally extending the milt and creating a usable and effective sperm solution for *in-vitro* fertilization or biobanking efforts.

Sperm quality varied between hormone treatments which could be related to the timing of gonadotropin release and subsequent LH and FSH function for maturation and provision of nutrients to sperm cells. It is unclear why the High treatment resulted in sperm with lower progressive and total motility; however, it is possible the GnRH-mediated gonadotropin release may become suppressed after receiving such high doses of GnRH, which could affect sperm production and function [43]. Alternatively, increased dopamine levels at higher doses of GnRH could be creating an antagonistic response downregulating the release of gonadotropins at the pituitary [44]. We observed that while resolving doses of GnRH past the Prime-Only did not affect concentration, the sperm that were released were of higher quality. Specifically, the Prime-Only treatment caused a release of sperm that were markedly lower in normal morphology compared to the other treatments possibly due to the clearing out of old dead cells. Even though the High treatment resulted in sperm samples with the lowest motility, it also resulted in the highest viability indicating that the cell membranes were protected. This difference could be due to GnRH having downstream effects on seminal fluid characteristics such as osmolality and ionicity that affect motility [45]. We observed that while further injections of GnRH past the Prime-Only did not affect concentration, it did seem to improve sperm quality. For example, the Prime-Only treatment had significantly lower numbers of viable sperm compared to the Medium and High treatments. There was a positive relationship between amount of hormone given and sperm viability, which could possibly be explained by differences in spermatogenesis and higher quality sperm being released at these higher hormone doses. The higher sperm abnormalities and lower viability of Prime-Only sperm samples could be due to older sperm cells being released which has been linked with elevated reactive oxygen species that can cause sperm cell membrane damage [46].

Our case study supports the transferability of ARTs protocols between salamander species. We found that GnRH treatment of 0.25 + 1.0 µg/g BW was able to elicit a spermic response from three out of four male arboreal salamanders tested. Moreover, this species also exhibited an episodic release of sperm, along with a wide range and variability in sperm concentration, that was similar to what was observed in the tiger salamanders. Although this is a limited sample size, it shows that small-bodied salamander species can be administered hormone in prey items with relative ease resulting in sperm release at levels applicable for biobanking or *in-vitro* fertilization purposes.

There were several limitations to this study including the number of study individuals. The few observed differences between treatments can likely be attributed to random and individual variation, and changes in peak sperm concentration times could likely differ. To note, one male released weakly concentrated spermic urine before receiving hormone but was considered to have a spermic response due to an increase in sperm concentration of two orders of magnitude after treatment. During one of the hormone rounds, *Acheta domesticus* crickets were substituted for *G. sigillatus* crickets due to logistical constraints, though their nearly identical size and structure should not have caused major differences in the overall response. The Prime-Only treatment round occurred last after the other three rounds of hormone, which could possibly be leading to decreased hormone sensitivity or sperm depletion within the animals; however, at least four weeks between hormone treatments has been found to be sufficient for generating independent responses [26,34]. Additionally, sperm was not collected until after the resolving dose, and spermiation in the first 24 hours is possible. However, previous work setting up hormone regimens for tiger salamanders collected sperm over the first 24 hours after just a prime dose and found that it took about 18 hours until the first milt was produced [26]. Tiger salamanders do have the ability to passively lay spermatophores when provided substrate [26], but we did not provide any, and no spermatophores were observed. Given the previous research with tiger salamanders that showed delayed sperm production with just an intramuscular prime, we decided to wait to collect until after the resolving dose.

The oral administration of hormone via a cricket vehicle not only improves the welfare of amphibians undergoing ART procedures by making the experience a positive event rather than a negative one, it also makes the technique available to a wider audience of practitioners such as husbandry staff [23]. Currently, most hormone administration through injection is restricted to veterinarians, academic researchers, curators, and trained scientists in zoos. While very promising, additional studies should examine the potential loss of hormone that could result from regurgitation or possible interactions with digestive enzymes within the salamander. Further expansion of oral hormone administration via prey items such as crickets should be applied to stimulate both natural and artificial breeding for small-bodied, at-risk caudates where ARTs were previously not feasible.

**S1 Data. Sperm data from oral hormone administration used for analysis.**

## Acknowledgements

Thank you to fellow members of the Mississippi State University Conservation Physiology Lab for their care of the animals that were part of this project.

